# Molecular characterization of carbapenem-resistant *Klebsiella pneumoniae* isolates with focus on antimicrobial resistance

**DOI:** 10.1101/778795

**Authors:** Xiaoling Yu, Wen Zhang, Zhiping Zhao, Chengsong Ye, Shuyan Zhou, Shaogui Wu, Lifen Han, Zhaofang Han, Hanhui Ye

**Affiliations:** Department of Infectious Diseases, Mengchao Hepatobiliary Hospital of Fujian Medical University, Xihong Road 312, Fuzhou 350025, Fujian, P.R. China; Friedman Brain Institute, Icahn School of Medicine at Mount Sinai, New York, NY 10029, USA; Key Lab of Urban Environment and Health, Institute of Urban Environment, Chinese Academy of Sciences, Xiamen 361021, Fujian, P.R. China; Department of Microbiology, Mengchao Hepatobiliary Hospital of Fujian Medical University, Xihong Road 312, Fuzhou 350025, Fujian, P.R. China; State Key Laboratory of Marine Environmental Science, College of Ocean and Earth Sciences, Xiamen University, Xiamen, 361102, Fujian, PR China; Xiamen Cingene Science and Technology co., LTD, Xiamen 361021, Fujian, PR China

**Keywords:** *Klebsiella pneumoniae*, Carbapenem-resistant, Prophage, whole-genome sequencing

## Abstract

The enhancing incidence of carbapenem-resistant *Klebsiella pneumoniae* (CRKP)-mediated infections in Mengchao Hepatobiliary Hospital of Fujian Medical University in 2017 is the motivation behind this investigation to study gene phenotypes and resistance-associated genes of emergence regarding the CRKP strains. In current study, seven inpatients are enrolled in the hospital with complete treatments. The carbapenem-resistant *K. pneumoniae* whole genome is sequenced using MiSeq short-read and Oxford Nanopore long-read sequencing technology. Prophages are identified to assess genetic diversity within CRKP genomes. The investigation encompassed eight CRKP strains that collected from the patients enrolled as well as the environment, which illustrate that *bla*_KPC-2_ is responsible for phenotypic resistance in six CRKP strains that *K. pneumoniae* sequence type (ST11) is informed. The plasmid with IncR, ColRNAI and pMLST type with IncF[F33:A-:B-] co-exist in all ST11 with _KPC-2_-producing CRKP strains. Along with carbapenemases, all *K. pneumoniae* strains harbor two or three extended spectrum β-lactamase (ESBL)-producing genes. *fosA* gene is detected amongst all the CRKP strains. The single nucleotide polymorphisms (SNP) markers are indicated and validated among all CRKP strains, providing valuable clues for distinguishing carbapenem-resistant strains from conventional *K. pneumoniae*. In conclusion, ST11 is the main CRKP type, and *bla*_KPC-2_ is the dominant carbapenemase gene harbored by clinical CRKP isolates from current investigations.

## Introduction

Antibiotic resistance is amongst the extremely severe public health challenges nowadays. Carbapenem-resistant *Enterobacteriaceae* (CRE) is reported as a consequence mainly due to acquisition of carbapenemase genes, and CRE is inferred as an urgent threat to human health by the Centers for Disease Control and Prevention (CDC), USA in 2013^1^. Carbapenems such as imipenem, meropenem, and biapenem represent the first-line treatment of serious infections caused by multi-resistant Enterobacteriaceae including *Klebsiella pneumoniae* (*K. pneumoniae*) and *Escherichia coli* (*E. coli*)^2^. Whereas carbapenems can be hydrolyzed by carbapenemase in carbapenem-resistant *K. pneumoniae* (CRKP)^3^, which results in resistance to β-Lactam antibiotics including carbapenem. Carbapenemases can be divided into Ambler class A β-lactamases (e.g. *Klebsiella pneumoniae* carbapenemases (KPC)), class B metallo-β-lactamases (MBLs), verona integrin-encoded metallo-β-lactamase (VIM), New Delhi metallo-β-lactamase (NDM) type, and Class D Enzymes of the OXA-48 type^4^. Among Ambler class A β-lactamases, plasmid-mediated KPC has been identified in all gram-negative members of the ESKAPE pathogens^5^, and KPC is the most clinically indispensable enzyme due to its prevalence in Enterobacteriaceae^6^. Moreover, pathogens harboring _KPC-2_ are resistant to all β-lactams and β-lactamase inhibitors except ceftazidime/avibactam, which extremely limit treatment options as well as lead to high mortality rates^7^. Additionally, NDM has become a serious threat to public health due to the rapid global dissemination of NDM-bearing pathogens and the presence on mobile genetic elements in an extensive series of species^8^. Consequently, it is imperative and urgent to investigate the CRKP characteristics for better controlling pathogens and diagnosing as well as treating patients.

In current investigation, seven CRKP strains are extracted from patients during their hospitalizations and another one CRKP strain is obtained from the dining car in Mengchao Hepatobiliary Hospital of Fujian Medical University (**Supplementary Table 1**). The whole genome of CRKP is sequenced using MiSeq short-reads and Oxford Nanopore long-reads sequencing technology. We conduct surveillance of the CRKP-mediated infection prevalence in Mengchao Hepatobiliary Hospital of Fujian Medical University, investigate the molecular epidemiological characteristics of the strains that obtained, and identify gene phenotypes as well as resistance-associated genes of the strain emergence. The detected single nucleotide polymorphisms (SNP) markers would be helpful for recognizing CRKP strain from general *K. pneumoniae*. Data of this study provide essential insights into effective strategy developments for controlling CRKP and nosocomial infection reductions.

## Materials & Methods

### Patient clinical information

In total, seven patients received treatments during their hospitalizations and the data of them were completely classified and studied. One bacterium was extracted from the dining car in the hospital and since the carrier was not human, there was no clinical data relating to it. All patients, except patient 1567P that was diagnosed as abdominal infection, were diagnosed as severe pneumonia or sufferred lung infections (**Supplementary Table 1**). We further give **Supplementary Table 2-8** to in detail provide all patients’ treatment records as well as the phenotype measurement results and data.

All patients received systematic medical examinations such as whole blood cell test, blood routine test, blood electrolyte test, blood clotting, fungal D-glucan detection, galactomannan detection, etc. All the records are archived in detail for further investigations.

### Bacterial Isolates and Antimicrobial Resistance

Single patient isolates are obtained from specimens that received from inpatients admitted to Mengchao Hepatobiliary Hospital of Fujian Medical University in 2017. A total of eight CRKP isolates (**Supplementary Table 1**), which are resistance to all the antibiotics tested, such as cephalosporins, penicilins, quinolones, aminoglycosides and carbapenems (Imipenem with MICs ≥16 μg/ml) (Table 1), were processed following standard operating procedures: the isolates are extracted according to the aseptic operating procedures and cultured in the bacterial culture medium with Columbia Agar + 5% sheep blood. The study has been performed in accordance with the Institutional Ethical Committee of the Faculty of Medicine, Mengchao Hepatobiliary Hospital of Fujian Medical University, which approved this study.

**Table 1.**
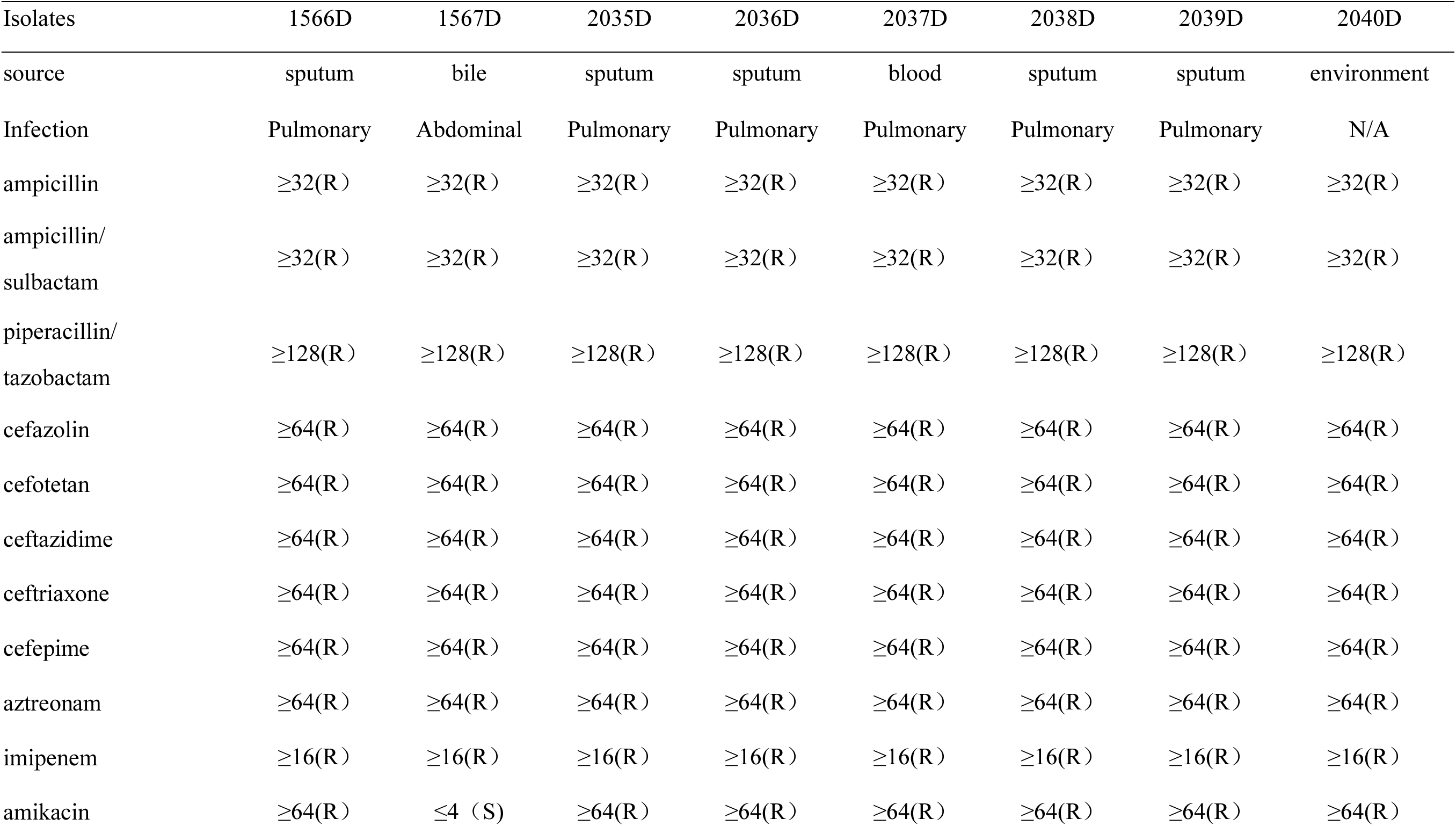
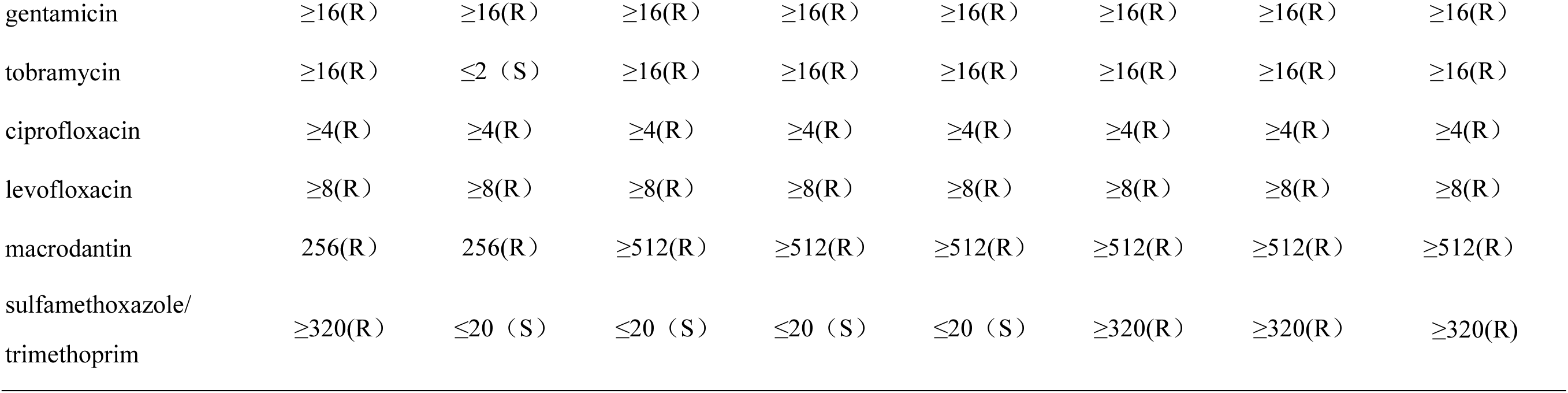
Antibiotic susceptibility profiles of *K. pneumoniae*. The results of antimicrobial susceptibility testing - antibiotics MIC (mg/L) and breakpoint interpretation or epidemiological cut-off value. S: susceptible; I: intermediate; R: resistant.

| Isolates | 1566D | 1567D | 2035D | 2036D | 2037D | 2038D | 2039D | 2040D |
| --- | --- | --- | --- | --- | --- | --- | --- | --- |
| source | sputum | bile | sputum | sputum | blood | sputum | sputum | environment |
| Infection | Pulmonary | Abdominal | Pulmonary | Pulmonary | Pulmonary | Pulmonary | Pulmonary | N/A |
| ampicillin | ≥32(R) | ≥32(R) | ≥32(R) | ≥32(R) | ≥32(R) | ≥32(R) | ≥32(R) | ≥32(R) |
| ampicillin/<br>sulbactam | ≥32(R) | ≥32(R) | ≥32(R) | ≥32(R) | ≥32(R) | ≥32(R) | ≥32(R) | ≥32(R) |
| piperacillin/<br>tazobactam | ≥128(R) | ≥128(R) | ≥128(R) | ≥128(R) | ≥128(R) | ≥128(R) | ≥128(R) | ≥128(R) |
| cefazolin | ≥64(R) | ≥64(R) | ≥64(R) | ≥64(R) | ≥64(R) | ≥64(R) | ≥64(R) | ≥64(R) |
| cefotetan | ≥64(R) | ≥64(R) | ≥64(R) | ≥64(R) | ≥64(R) | ≥64(R) | ≥64(R) | ≥64(R) |
| ceftazidime | ≥64(R) | ≥64(R) | ≥64(R) | ≥64(R) | ≥64(R) | ≥64(R) | ≥64(R) | ≥64(R) |
| ceftriaxone | ≥64(R) | ≥64(R) | ≥64(R) | ≥64(R) | ≥64(R) | ≥64(R) | ≥64(R) | ≥64(R) |
| cefepime | ≥64(R) | ≥64(R) | ≥64(R) | ≥64(R) | ≥64(R) | ≥64(R) | ≥64(R) | ≥64(R) |
| aztreonam | ≥64(R) | ≥64(R) | ≥64(R) | ≥64(R) | ≥64(R) | ≥64(R) | ≥64(R) | ≥64(R) |
| imipenem | ≥16(R) | ≥16(R) | ≥16(R) | ≥16(R) | ≥16(R) | ≥16(R) | ≥16(R) | ≥16(R) |
| amikacin | ≥64(R) | ≤4 (S) | ≥64(R) | ≥64(R) | ≥64(R) | ≥64(R) | ≥64(R) | ≥64(R) |
| gentamicin | ≥16(R) | ≥16(R) | ≥16(R) | ≥16(R) | ≥16(R) | ≥16(R) | ≥16(R) | ≥16(R) |
| tobramycin | ≥16(R) | ≤2 (S) | ≥16(R) | ≥16(R) | ≥16(R) | ≥16(R) | ≥16(R) | ≥16(R) |
| ciprofloxacin | ≥4(R) | ≥4(R) | ≥4(R) | ≥4(R) | ≥4(R) | ≥4(R) | ≥4(R) | ≥4(R) |
| levofloxacin | ≥8(R) | ≥8(R) | ≥8(R) | ≥8(R) | ≥8(R) | ≥8(R) | ≥8(R) | ≥8(R) |
| macrodantin | 256(R) | 256(R) | ≥512(R) | ≥512(R) | ≥512(R) | ≥512(R) | ≥512(R) | ≥512(R) |
| sulfamethoxazole/<br>trimethoprim | ≥320(R) | ≤20 (S) | ≤20 (S) | ≤20 (S) | ≤20 (S) | ≥320(R) | ≥320(R) | ≥320(R) |

*K. pneumoniae* isolates are confirmed by Matrix-assisted Laser Desorption Ionization-time of Flight Mass Spectrometry (MALDI-TOF-MS) mass spectrometry (BioMerieux SA, BioMerieux Inc., France). The resistance of pathogenic bacteria is identified by Automatic Microbial Identification & Drug Sensitivity Analysis System (VITEK-2 Compact, BioMerieuxInc., France) with Gram-Negative identification card (VITEK2 AST-GN13, BioMerieuxInc., France). The results of antimicrobial susceptibility testing are interpreted based upon Clinical and Laboratory Standards Institute (CLSI) M100-S24^9^. The standard strain under quality control is *K. pneumoniae* isolates ATCC700603 (American Type Culture Collection, ATCC).

### Whole Genome Sequencing (WGS) and Assembly

The isolated seven CRKP bacteria are sequenced on Illumina MiSeq (Illumina, San Diego, CA, USA) platform. MiSeq short-read sequencing library is generated with 1 ng purified DNA. Inserting a phosphate to 5’ UTR end and “A” to 3’ UTR end produces end-repair, and PCR fragments (300 ∼ 600 bp) are collected from bar-coded adapter ligation. The library is purified via AMPure XP (Beckman Coulter), which is then sequenced on MiSeq platform. In sum, a total of 40.5 million reads (2 × 300 bp) with a size of 1.36 Gb data are yielded (**Supplementary Table 9**). All short reads are first filtered for the low-quality sequences and then assembled into contigs using SPAdesv3.11.1 software^10^.

Subsequently, we select an isolate of 1567D to perform long-read sequencing on Oxford Nanopore MinION (Oxford, UK) platform to easily sequence across repeat regions. The sequencing library is constructed with 1.5 μg purified DNA using the LSK-108 Oxford Nanopore Technologies (ONT) ligation protocol, and the prepared library is sequenced following the standard protocol of Oxford Nanopore MinION. A total of 7.48 Gb ultra-long reads are generated with N50 length of 25,890 bp (**Supplementary Table 10**). The long reads that ‘passed’ during the Nanopore base calling are used to assemble into complete genomic sequences via Canu software^11^. The long-read sequencing data of the same individual are used to correct base errors of assembled genome using Nanopolish (https://github.com/jts/nanopolish).

### Detecting Prophages in the CRKP Genomes

The putative prophages within contigs of the CRKP genomic sequences are identified using the PHAST web server (PHAge Search Tool)^12^. The prophage completeness and categorization (intact, incomplete, or questionable) are presented applying over sequences to check homology, and to detect, annotate, and graphically display prophages.

### Carbapenemase-resistance Gene Identifications

To predict the protein-coding genes and functional proteins in the CRKP genomes, all assembled sequences are annotated by a web-based package RAST (Rapid Annotations using Subsystems Technology)^13^. The antibiotic resistance and virulence genes, plasmids, phenotyping and genotyping of CRKP genomes are scanned using the Bacterial Analysis Pipeline^14^. Carbapenemase-resistance genes are further identified from above annotated sequences according to Simner et al.^15^.

The protein-coding genes of long-read assembled genome are predicted using GLIMMER (Gene Locator and Interpolated Markov ModelER) v3.02^16^. To functionally annotate the predicted genes and perform the pathway analysis, we align them to NR, COG, Swiss-Prot, GO and KEGG databases using blastX (E-value: 10^−5^). The annotated genes serve to improve the completeness of some important carbapenemase-resistance genes.

Comparisons of strain similarity are performed using the Harvest Tools Suite^17^ (version 1.1.2). For all of the isolates sequenced on a particular platform, parsnp is utilized to compare all the assembled isolates against each other and known reference strain. Results are visualized using EvolView.

### SNP Identification and Validation

We download *K. pneumoniae* genome from NCBI as the reference to identify SNP markers^18^. All high-quality data (Q value >20, reads length > 50 bp, number of uncertain bases < 5%) of eight CRKP strains are aligned to the reference genome sequences using BWA v0.7.17^19^, and aligned reads are sorted by coordinates via SAMTOOLS v1.4^20^. The GATK (Genome Analysis Tool Kit) software v3.8.0^21^ is utilized to detect SNPs, which is described as following: (1) duplicated reads are removed; (2) reads around insertions/deletions are realigned; (3) base quality is recalibrated using default parameters; (4) all variants are identified using HaplotypeCaller method in GATK with emitting and calling standard confidence thresholds at 10.0 and 30.0, respectively. To validate the detected SNPs in the seven CRKPs, we select 20 loci within each sample that are located in exonic regions and sequence them with high read depth. All chosen markers are designed primers for amplification using Sequenom MassARRAY iPLEX platform.

## Results

### Antimicrobial Susceptibilities of the CRKP Strains

The source of isolates is supplied in **Table 1**, which denotes the infectious type and the result of susceptibility testing during the patients’ hospitalization. All eight strains involved in the study are confirmed to be *K. pneumoniae*, with five strains from sputum, one from bile, one from blood, and one from the environment (**Supplementary Table 1**). Clinical data demonstrate that seven of the eight patients are referred due to pulmonary infection, and another one is referred due to abdominal infection. The susceptibility testing data in **Table 1** reveals that all the *K. pneumoniae* strains are resistant to almost all antibiotics, such as cephalosporins, penicilins, quinolones and carbapenems (imipenem with MICs ≥16 μg/ml). For aminoglycosides antibiotics, except that 1567D isolate is sensitive to amikacin and tobramycin, all other isolates are resistant to aminoglycosides antibiotics. The strains including 1566D, 2038D, 2039D and 2040D are resistant to sulfamethoxazole/trimethoprim with MICs ≥320, and the other strains (1567D, 2035D, 2036D, 2037D) are sensitive to sulfamethoxazole/trimethoprim with MICs ≤20.

### Genome assembly and annotation

The short-read sequenced seven CRKP strains are assembled into contigs. As listed in **Table 2**, the assembled genome size of all trains ranged from 5.4 Mb to 5.8 Mb, with mean length of 5.7 Mb and average contigs numbering 199. The N50 length of genomes is from 176.6 kb to 251.6 kb with an average N50 length of 220.4 kb and mean GC content of 57.2%. To obtain a more complete genome, the 1567D strain is resequenced via long-read sequencing technology and assembled into three contigs with size of 5.6 Mb (**Supplementary Figure 1**). A total of 5,841 protein-coding genes are predicted with length between 37 to 1,649 bp (**Supplementary Figure 2**). Totals of 4,657, 5,097, 4,714, 3,179 and 3,099 predicted genes are functionally annotated in NR, COG, Swiss-Prot, GO and KEGG databases, respectively (**Supplementary Figures 3, 4, 5**).

**Table 2.** Assembly statistics of seven CRKP strains via short-read sequencing.

| Assembly | 1566D | 2035D | 2036D | 2037D | 2038D | 2039D | 2040D |
| --- | --- | --- | --- | --- | --- | --- | --- |
| Contig number | 195 | 163 | 215 | 242 | 184 | 196 | 195 |
| Total length (bp) | 5,758,754 | 5,633,502 | 5,432,179 | 5,670,795 | 5,833,697 | 5,831,354 | 5,835,044 |
| Largest contig (bp) | 380,781 | 381,132 | 929,110 | 380,795 | 381,132 | 381,056 | 381,056 |
| GC (%) | 57.33 | 57.39 | 57.18 | 57.35 | 57.21 | 57.21 | 57.2 |
| N50 | 176,606 | 196,479 | 251,620 | 196,479 | 183,862 | 190,289 | 183,862 |
| L50 | 12 | 11 | 7 | 11 | 11 | 11 | 11 |
| Total number of Ns | 20 | 30 | 20 | 30 | 130 | 130 | 30 |

### Characteristics of the CRKP Isolates

The isolated seven CRKP bacteria are sequenced through Illumina MiSeq platform and assembled into whole genomes. To understand genetic diversity, mobile genetic elements of 24 prophages are identified in seven CRKP genomes, with sizes ranging from 8.4 kb to 49.3 kb (**Figure 1**). According to the criterion that the length of an intact prophage should be more than 20 kb^22^. Prophages detected in most strains (except for 2036D) are complete with a size of at least 20.2 kb with an average GC percentage of 52.7%. Additionally, three prophages are respectively identified in 3 strains at the same time, revealing the genomic sequence homology among all isolates. The 2036D strain is comprised of just one prophage probably because of the small genome size and distinct sequence characteristics, which is expected to have less neutral targets for prophage integration^22^.

**Figure 1.**
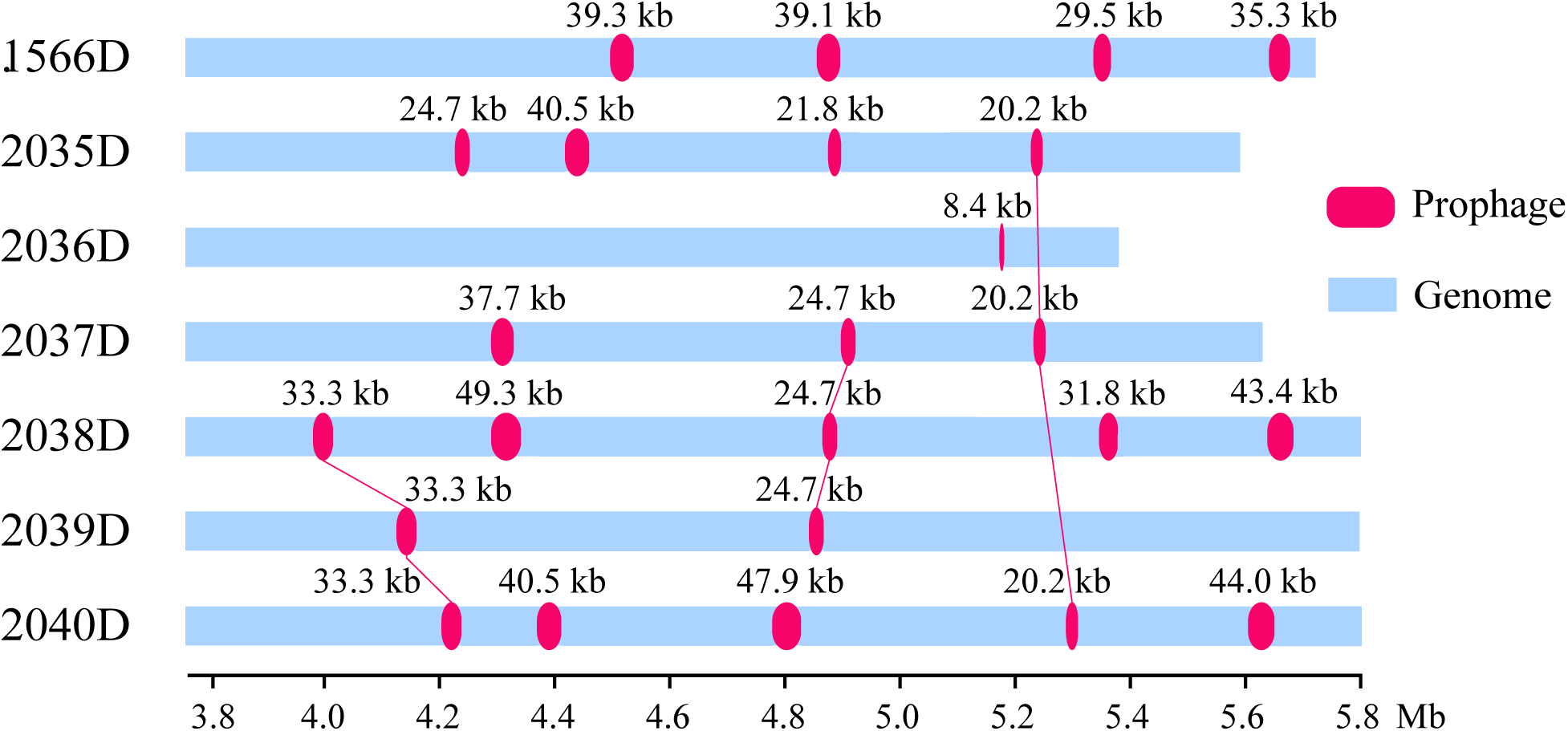
Intact prophages identified in seven CRKP strains.

Furthermore, multilocus-sequence typing (MLST) analysis reveals that there are two unrelated sequence type (ST) in *K. pneumoniae* strains isolated from different patients. 2036D *K. pneumoniae* strain correlates with ST2632, and the other six strains are relevant to ST11 (**Table 3**). pMLST analysis reveals that all of the six ST11 *K. pneumoniae* strains are associated with IncF[F33:A-:b-] and the ST2632 *K. pneumoniae* strain is relevant to IncHI1 and IncF. Majority of *K. pneumoniae* strains are ST11, with IncF [F33:A-:b-] type.

**Table 3.**
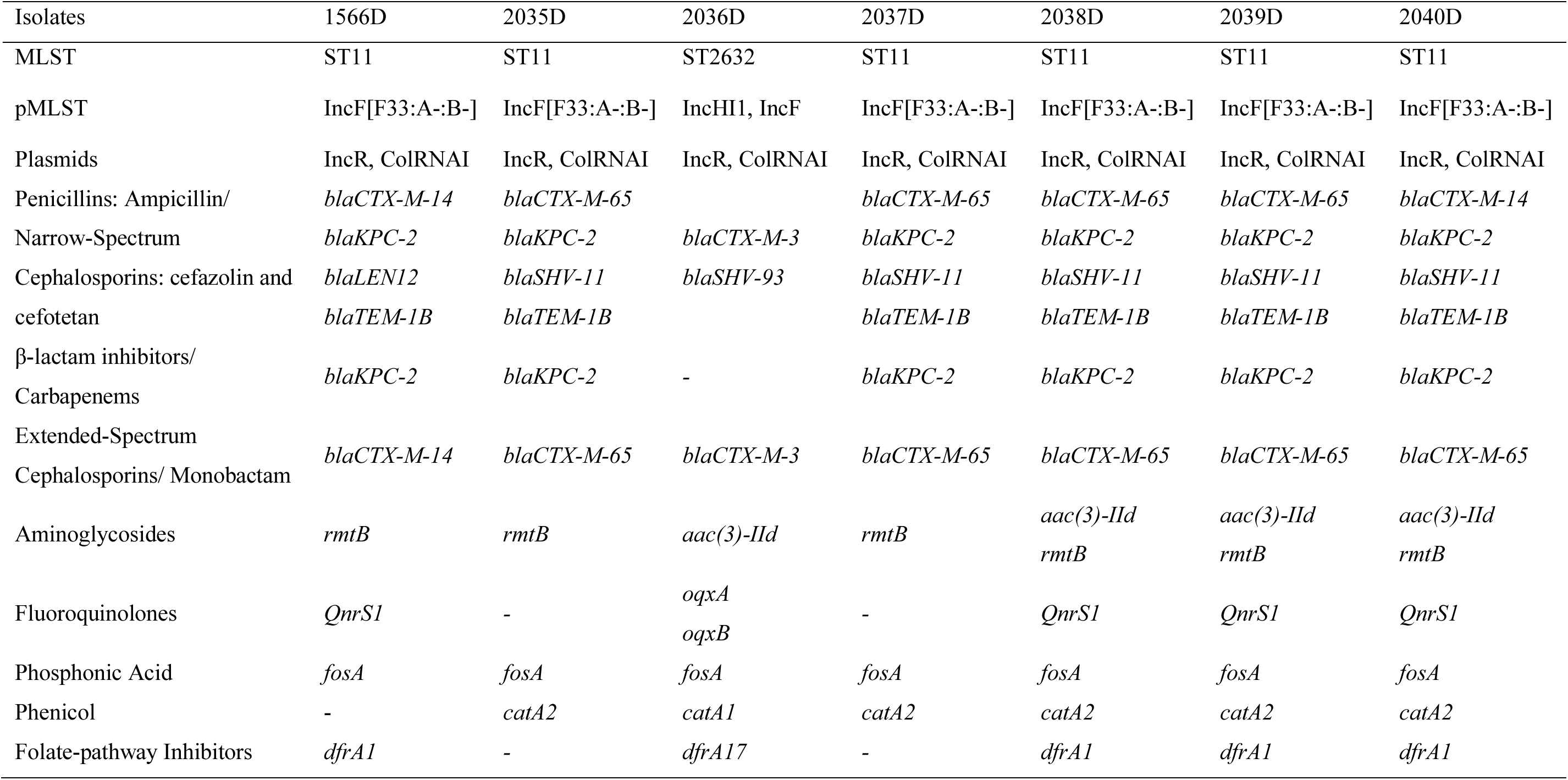
Resistance genes among the patient and environmental isolates.

Plasmid analysis^23^ shows different circular plasmids carried by the individual strains. All strains harbored IncR and ColRNAI plasmids with no virulence genes but contain several resistance-associated genes that cause resistance to carbapenems, which is demonstrated in **Table 3**. The IncR plasmid is identified as multidrug-resistant plasmids and has variable copy numbers of certain resistance genes among *K. pneumoniae* isolates.

### Detection of Antibiotic Resistance Genes of CRKP Isolates

The resistance-associated genes of seven CRKP bacteria (**Table 3**) are sequenced on Illumina MiSeq platform among the patient and environmental isolates. As illustrated in **Table 3**, some antimicrobial resistance genes are mediated by plasmid such as β-lactamase correlative genes (*bla*_CTX-M_, *bla*_KPC_, *bla*_LEN_, *bla*_TEM_) and those genes which encoded aminoglycoside [*aac(3)-IId, rmtB*], chloramphenicol (*catA1, catA2*), trimethoprim (*dfrA1,dfrA17*), and fluoroquinolone [*QnrS1*]. The other antimicrobial resistance genes are encoded by chromosome including *bla*_SHV_ (narrow-spectrum β*-*lactamasein *K. pneumoniae*), *oqxA* (1,176 bp), *oqxB* (3,153 bp) (efflux pumps), and *fosA* (420 bp, fosfomycin resistance) genes.

Except 2036D, all the other *K. pneumoniae* strains harbor the associated carbapenemases-producing resistance gene, *bla*_KPC-2_. Extended-spectrum β-lactamases (ESBLs) resistance genes such as *bla*_CTX-M_, *bla*_TEM_, *bla*_LEN_ and *bla*_SHV_ are also informed. *bla*_TEM_ is one of the genes that produce ESBL. *bla*_CTX-M_ with different types (*bla*_CTX-M-14_, *bla*_CTX-M-3_, *bla*_CTX-M-55_ and *bla*_CTX-M-65_) is found among all the *K. pneumoniae* strains. *bla*_CTX-M-3_ is observed in 2036D strains. *bla*_CTX-M-55_ is observed in 1567D strain and *bla*_CTX-M-_14 is observed in 1566D and 2040D strains. *bla*_CTX-M-65_ is detected in the other four (2035D, 2037D, 2038D, 2039D) *K. pneumoniae* strains. *bla*_LEN12_ gene is exclusively found in 1566D strain, and there is no *bla*_SHV_ gene in it. Nevertheless, *bla*_SHV-93_ and *bla*_SHV-11_ genes are detected in 2036D strain and the other five *K. pneumoniae* strains, respectively. Except for the 2036D strain, *bla*_TEM-1B_ gene is observed in all the other six *K. pneumoniae* strains. *Aac(3)-IId* and *rmtB* encoding fluoroquinolone resistance are observed among all strains. *fosA* resulting in fosfomycin resistance^24^ is also informed among all strains.

### Characterizing CRKP SNPs

The SNP markers are identified for all strains that sequenced using the short-read MiSeq data. The data demonstrate that 33,716 markers are detected in the 2036D strain, which is more significant than the other strains with an average of 8,289 SNPs. The cSNPs located in exonic regions are in slightly higher amounts among all detected SNPs of a minimal ratio of 85.5% (**Supplementary Table 11**). In addition, the pairwise comparison analysis reveals that 2036D isolate is disparate with the other strains based on clusters of sequence similarities using subprogram of Trinity^25^ (**Figure 2a**). Furthermore, the 2036D strain share few SNP loci with the others, which coincides with strain clusters (**Figure 2b**).

**Figure 2.**
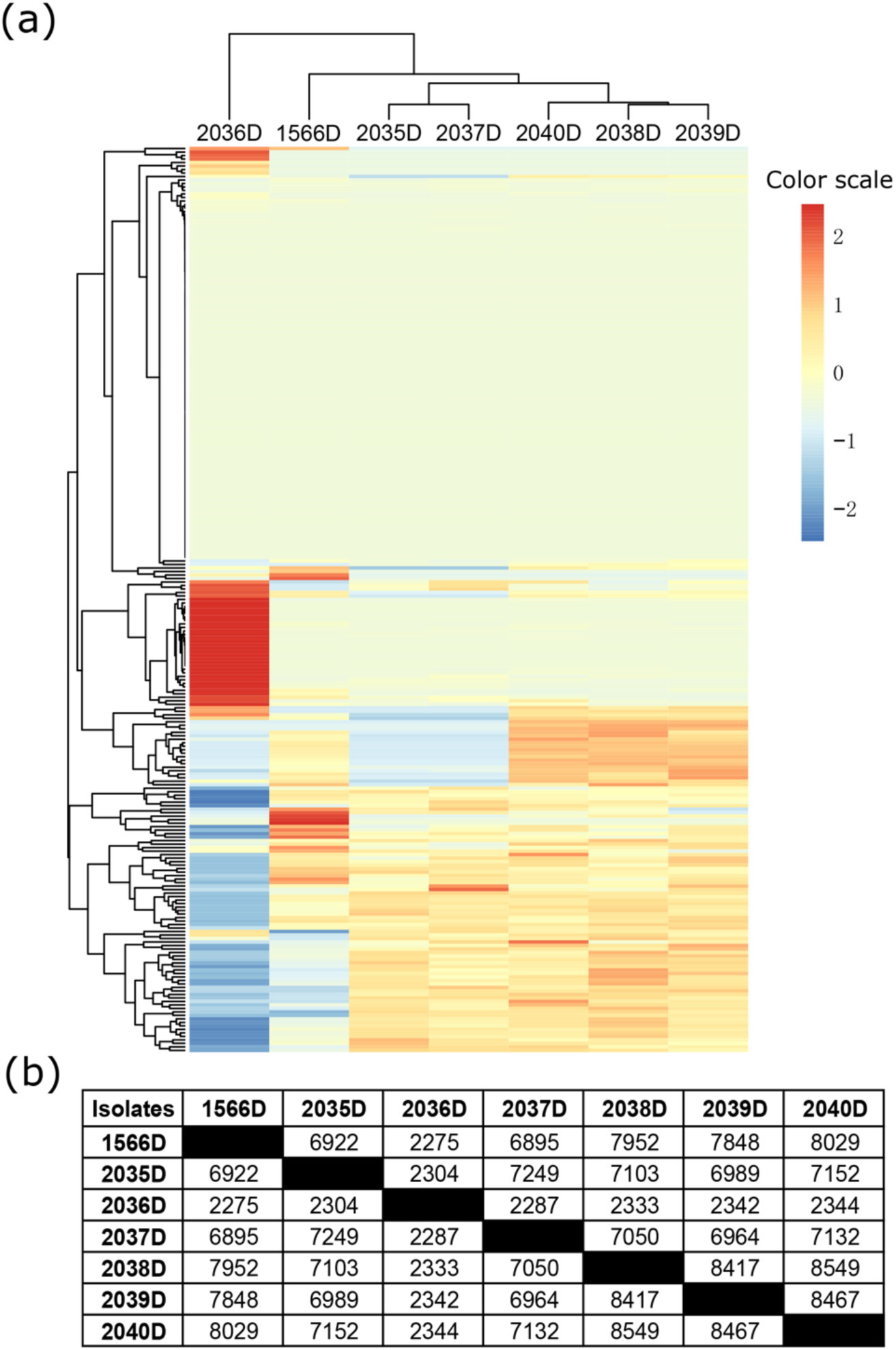
Assessing the genetic relatedness of the CRKP by WGS. (a) Clustering of all isolates based on sequence similarity. (b) Communal SNP markers detected by pairwise comparison analysis.

For validations, all strains have a high detection rate in that approximately 153 out of 200 SNPs (76.4%) that have amplifications, which demonstrate the analysis accuracy (**Supplementary Table 11**). After filtering SNP loci that are not located in exome regions, containing no-alleles locus, and comprise all-wild SNP loci in each isolate, we eventually obtain 92 SNPs among 200 validated loci. A total of 40 out of 92 SNPs are all-variation loci in all isolates, which could be utilized for recognizing CRKP strain from ordinary *K. pneumoniae* (**Supplementary Table 12**). In addition, 24 SNPs of strain’s unique loci, including strains of 2036D (18 loci), 2035D (3 loci), 1566D (2 loci) and 2037D (1 loci), would be helpful resources for specific strain identification of clinical analysis.

### Phylogeny analysis results

The phylogenic tree shows that 1567D strain is most distantly related to the other strains, and 2036D is more closely related to the reference strain comparing with all the other strains (**Supplementary Figure 6**).

### GWAS analysis

To further identify significant SNPs and genes, we perform genome-wide association study (GWAS) analysis. The patients’ body temperature and counts of leukocyte are selected as phenotypic character. The short-sequencing reads of six strains (**Figure 3**) are aligned to the 1567D genome using BWA v0.7.17 software. We call SNPs using Platypus v0.8.1^26^, and then filter the SNPs through plink v1.9 according to the following conditions: (i) missing loci, (ii) minor allele frequency (MAF) < 0.05 and (iii) significant deviation from the Hardy-Weinberg equilibrium (HWE) (*P* < 0.01). A total of 698 SNP markers are remained and utilized for GWAS analysis. As a result, 9 loci are identified (*P* < 0.05). Two loci (ygbI and murB) are related with temperature and the other seven loci (IsrD, SufD, yrkF, fabI, sppA, entF and ttuB) are relevant to leukocyte (**Figure 3**).

**Figure 3.**
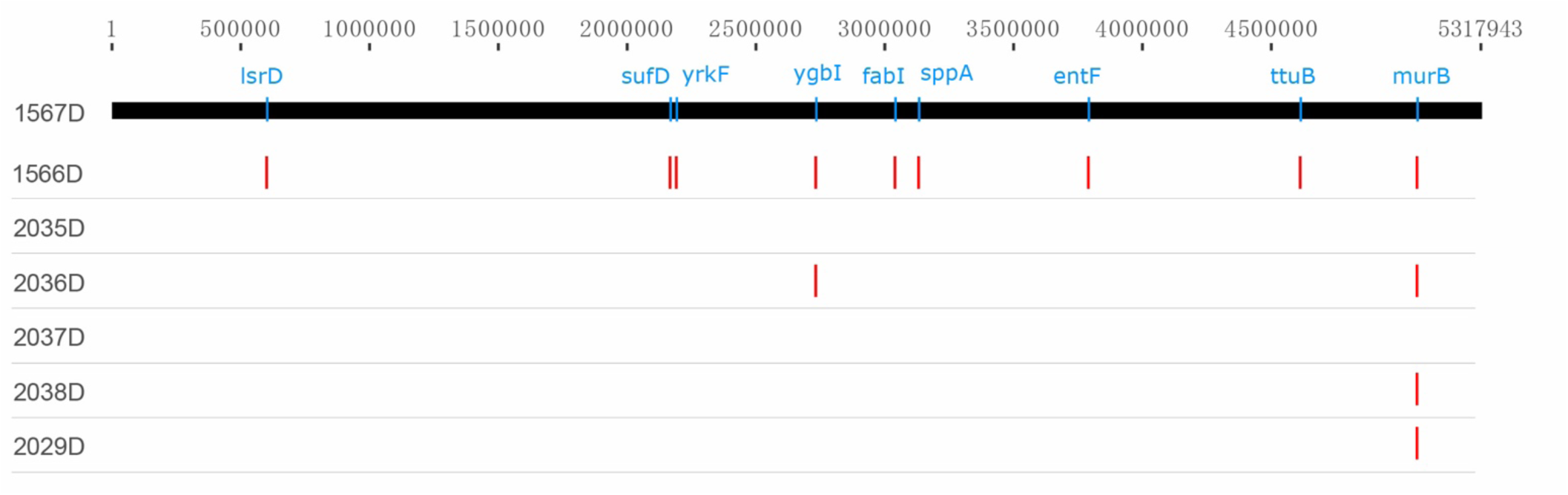
GWAS results of the analysis.

## Discussion

Data of current study confirm that all CRKP strains hold two types of plasmids with no virulence gene whereas harbor an abundance of associated resistance genes such as ESBLs and carbapenemases. One genotype of carbapenemases with *bla*_KPC-2_ and two ST types with ST11 and ST2632 are identified in the study, and the ST11 with KPC-2-positive is a prevalent strain accounting in all the six strains. The plasmid with IncR, ColRNAI and pMLST type with IncF[F33:A-:B-] co-exist in all ST11 with KPC-2-producing CRKP strains. The initial detection of a KPC-2-producing *K. pneumoniae* isolate from a hospital in China is reported in 2007^27^. Since then, *bla*_KPC-2_-bearing *K. pneumoniae* isolates have become more prevalent and reported in China as well as other countries and areas ^28^. Recently, one patient is found to have susceptible *K. pneumoniae* bacteraemia in US^15^. While that case is relatively specific since the patient might be affected during the visit and hospitalization in India, which would add more complex environmental factors to confound the results. CRKP of ST11 associated with *bla*_KPC-2_ is disseminated widely across China^18,29^, which is concordant with the results of our study. These findings all suggest that the CRKP-mediated infections in our hospital result from ST11 with KPC-2-positive *K. pneumoniae* isolates. Continuous monitoring will be necessary to prevent further dissemination of carbapenemase-resistance genes.

Besides carbapenemases, a variety of ESBLs such as *bla*_CTX-M_, *bla*_SHV_, *bla*_LEN_, *bla*_TEM_ are present in CRKP strains of this study. *K. pneumoniae* is one of the most indispensable infectious agents in the ICU^30^. There are “classic” and hypervirulent strains of *K. pneumoniae*^31,32,33^. The “classic” non-virulent strain of *K. pneumoniae* (C-KP) can produce ESBLs related to nosocomial infectious outbreaks especially in the ICU of a hospital. C-KP more easily acquires antimicrobial resistance such as ESBLs. *bla*_CTX-M_ with different type is found among all the CRKP strains. Chromosome-mediated *bla*_SHV_ and plasmid-mediated *bla*_TEM_ are also positive for ESBLs production and are observed in six *K. pneumoniae* strains. Co-occurrence of *bla*_CTX-M_, *bla*_KPC-2_, *bla*_SHV-11_ and *bla*_TEM-1B_ are observed among five *K. pneumoniae* strains. All *K. pneumoniae* strains harbor two or three ESBLs-producing genes (*bla*_CTX-M_, *bla*_SHV_ and *bla*_TEM_), which indicate all isolates contained multiple ESBLs resistance genes. Previous reports noted consistent results that co-occurrence of *bla*_TEM_, *bla*_SHV_ and *bla*_CTX-M_ (any two or all three) was observed among *Klebsiella* isolates^34^.

*fosA* is frequently identified in the *E. coli* and *K. pneumoniae* genomes ^35,36^. The *fosA5* gene is first found in *E. coli* in 2014^37^. In 2017, it was reported that all of 73 carbapenem-resistant *K. pneumoniae* isolates were positive for *fosA5* in one Chinese area: Zhejiang Province^38^. Antimicrobial susceptibility testing about fosfomycin is not conducted in this study though *fosA* is also found among all the CRKP strains, which might indicate that fosfomycin-modifying enzymes account for a majority of the fosfomycin resistance, and that fosfomycin is resistant to CRKP strains. It is reported that *fosA* gene is transferred from *E. coli* to *K. pneumoniae* through whole plasmid transmission or mobile genetic element transmission, which raise doubts whether fosfomycin can be used as a supplementary drug for urinary tract infection caused by carbapenem-resistant *E. coli* in the hospital, as *fosA* exists in all CRKP strains from our study. Continuous monitoring will be necessary to prevent further dissemination of fosfomycin-resistant bacteria together with prudent use of fosfomycin in clinical settings.

*OqxA* and *oqxB* genes are relevant to efflux pumps, which means that antibiotics such as cephalosporins, carbapenems and fluoroquinolones are almost completely expelled from *K. pneumoniae* through its cell membrane^39^. Although there are no carbapenemases that observed in 2036D strain, *oqxA* and *oqxB* genes are identified in it. To our knowledge, these two genes are mainly reported to be responsible for the resistance to fluoroquinolones. They do have been previously reported to be associated with the nitrofurantoin resistance.

The genome sequences of the seven strains include massive contigs which are highly fragmented. Upon further investigation, we sequence the 1567D strain using long-read sequencing platform, which could help us assemble the genome with considerable improvement in completeness and contiguity. The carbapenem-resistant genes including *fosA, oqxA* and *oqxB* and 40 all-variation SNP loci are also identified in the above genome demonstrating the high-quality assembly. The assembly and annotation information will be beneficial in understanding the whole genomic characterization of CRKP strain for future study.

## Conclusions

In conclusion, ST11 is the main CRKP type, and *bla*_KPC-2_ is the dominant carbapenemase gene harbored by clinical CRKP isolates from present study. The plasmid with IncR, ColRNAI and pMLST type with IncF[F33:A-:B-] exist in all ST11 with KPC-2-producing CRKP strains. Besides carbapenemases, all *K. pneumoniae* strains harbor two or three ESBLs-producing genes (*bla*_CTX-M_, *bla*_SHV_ and *bla*_TEM_), which indicate all isolates contain multiple ESBLs resistance genes. *fosA* genes are also found among all the CRKP strains, which may infer that fosfomycin-modifying enzymes account for a majority of the fosfomycin resistance and that CRKP strains are resistant to fosfomycin. The differential expressions of *oqxA* and *oqxB* in CRKP strain might possibly result in carbapenem-resistant, but this presumption needs more solid experimental evidences. The 40 all-variation SNP loci in all isolates could be employed and referred for distinguishing CRKP strain from ordinary *K. pneumoniae*.

## Declarations

### Ethics approval

The study has been performed in accordance with the Institutional Ethical Committee of the Faculty of Medicine, Mengchao Hepatobiliary Hospital of Fujian Medical University.

### Consent to participate

Not applicable.

### Consent to publish

All authors read and approved the final manuscript.

### Availability of data and materials

The genome shotgun sequencing data and long reads of Oxford Nanopore data are deposited at NCBI/GenBank as BioProject of PRJNA506754.

### Authors’ Contributions

X.Y., Z.Z., L.H. and H.Y. conceived of the method. H.Y. supervised the study. S.Z., S.W. and C.Y. implemented the bacteria culture. X.Y., W.Z. and Z.H. performed the bioinformatics analysis. X.Y. and Z.H. optimized and performed the sequencing. X.Y., W.Z. and Z.H. drafted the article with inputs and feedbacks from all the other authors. All authors read and approved the final manuscript.

## Acknowledgements

Authors express their gratitude’s to Rebecca Lahniche from the University of Pittsburgh English Language Institute for the proofreading assistance.

