## supplementary Information for "Molecular characterization of carbapenem-resistant *Klebsiella pneumoniae* isolates with focus on antimicrobial resistance"

**Supplementary Figures 1 to 6**
**Supplementary Tables 9 to 12** (**Supplementary Tables 1-8** are in separate Excel files)

### Supplementary Figures

**Supplementary Figure 1. Circle diagram of K. pneumoniae genome sequenced via Oxford Nanopore sequencing technology.**

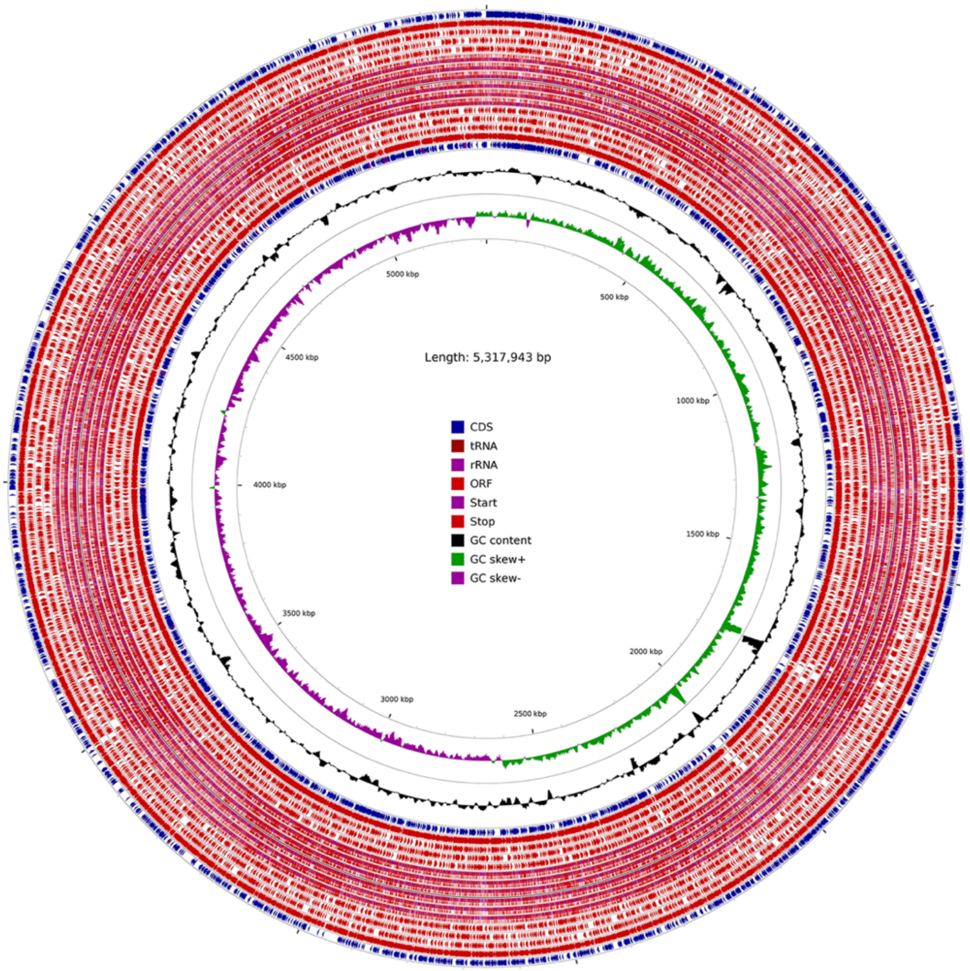

**Supplementary Figure 2. Distribution of protein-coding genes predicted in 1567D strain.**

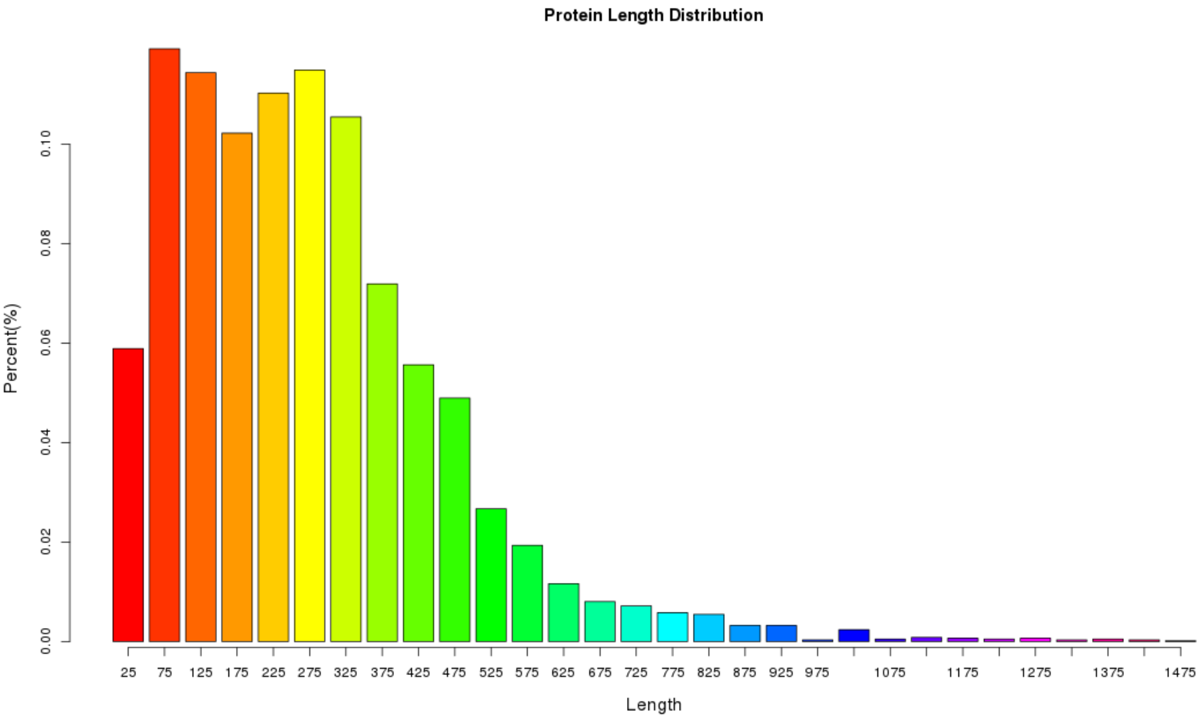

**Supplementary Figure 3. COG classification 1567D stain for the carbapenem-resistant K. pneumoniae.**

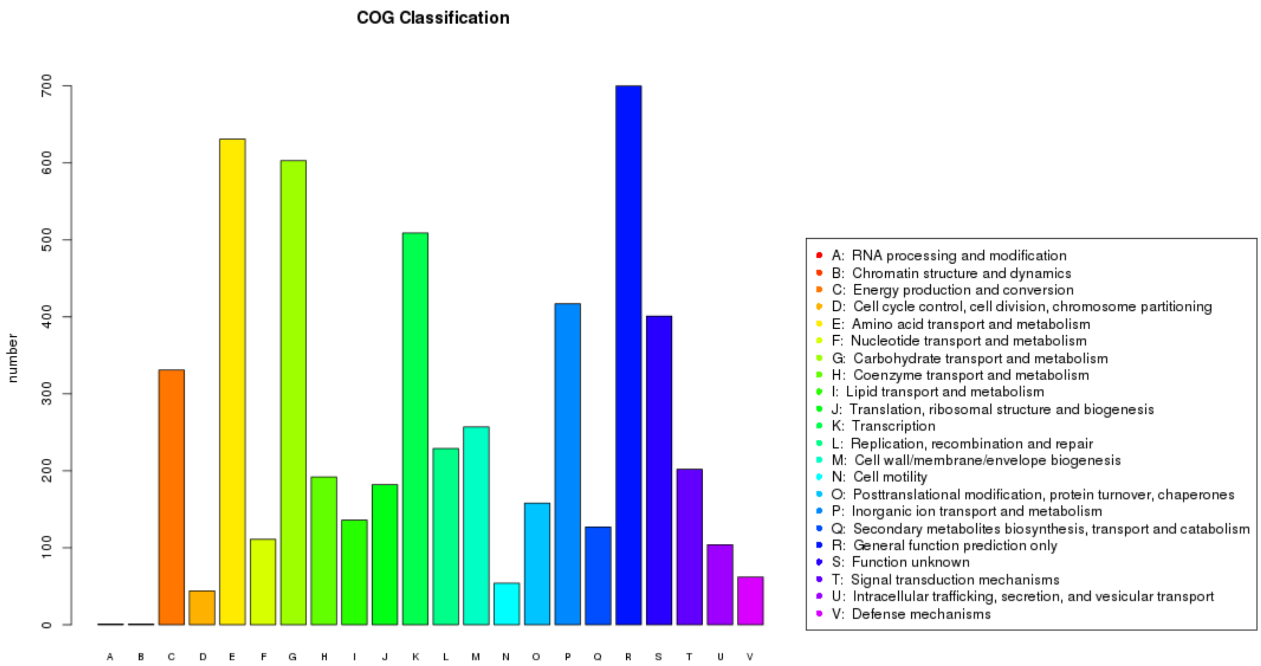

**Supplementary Figure 4. Distribution of K. pneumoniae genes annotated in GO term.**

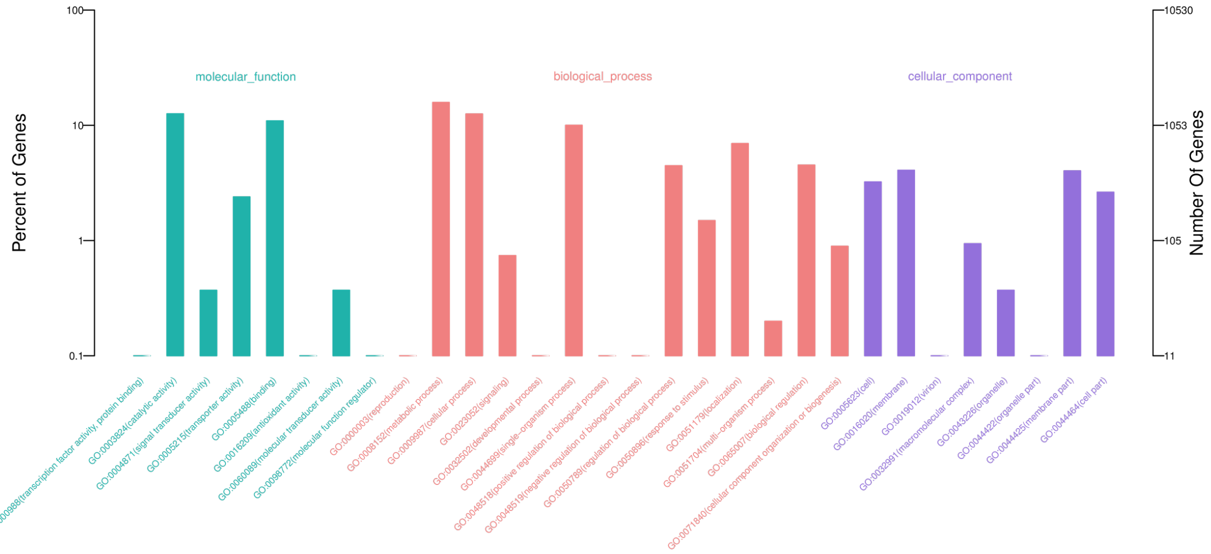

**Supplementary Figure 5. Annotation of KEGG pathways in the carbapenem-resistant K. pneumoniae.**

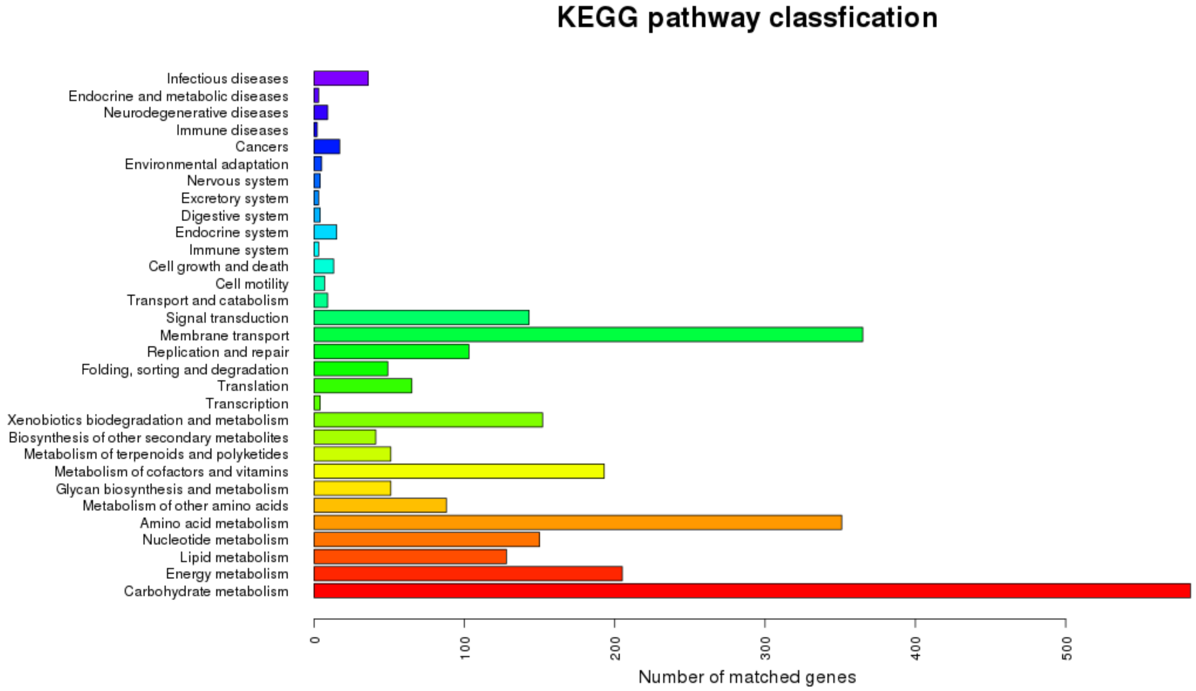

**Supplementary Figure 6. Phylogenetic tree assessing the relatedness of the 8 carbapenem-resistant *K. pneumoniae* strains (red) to the reference genome database (blue).**

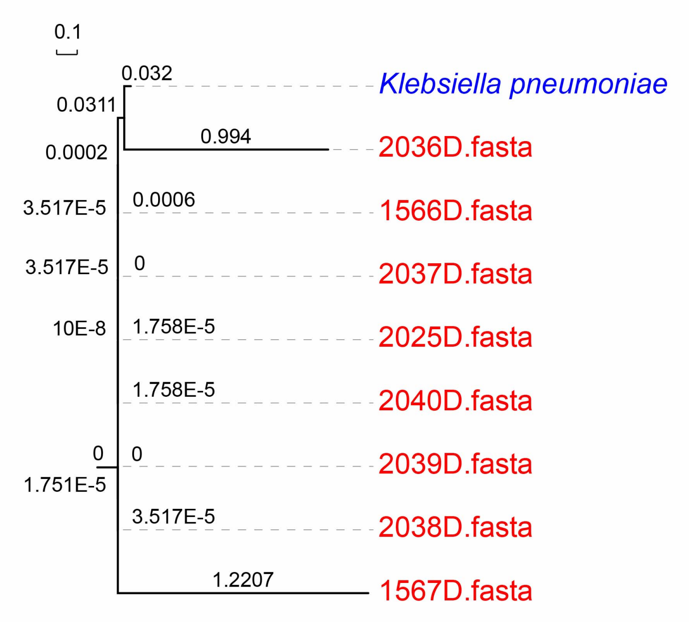

#### Supplementary Tables

All patients, except patient 1567P that was diagnosed as abdominal infection, were diagnosed as severe pneumonia or sufferred lung infections (**Supplementary Table 1**). We give **Supplementary Table 2-8** in detail to provide all patients' treatment records as well as the phenotype measurement results and data. For instance, 2036P (**Supplementary Table 5**) is an 83-year-old male patient. Previous to being admitted in our hospital, he was treated in another hospital. His medical record as of August 1st, 2017 showed that the urea nitrogen was 28.1 mmol/L, and creatinine was 429 µmol / L. His urinary red blood cell malformation rate was 56% high. Pulmonary CT indicate that there were nodules in his right upper lung, possibly being peripheral lung cancer. There were bilateral pleural effusion. He was hospitalized in our hospital from August 25th to September 29th, 2017. For his and all other patients' medical treatment records, please refer to **Supplementary Table 2-8**.

All the eight strains (including the environmental isolate 2040D) were collected from the same floor (within the same ward) in our hospital (Supplementary Table 1). Since the first CRKP (2035D) was reported on October 21, 2017, other CRKP's were detected intermittently in October (2036D), November (2037D), and December (2038D and 2039D), 2017 in our hospital. It caused our attentions and we took interventions accordingly. After the above cases were reported, the microbiology room retained all the resistant strains since the hospital had a routine drug resistance monitoring. The CRKP's that detected before October 21, 2017 were tracked and extracted from the strain bank, which were resuscitated. We did sequencing analysis for the additional two CRKP's (1566D and 1567 D) together with the above CRKP's. We analyzed them together in the article.

We collected the dining car that held lunches and dinners of the patients on that ward. We detected hands of the medical staff as well, and finally found the CRKP (2040D) on the dining car. No CRKP were detected elsewhere. With this source, the purpose of sequencing the bacteria car is to validate whether it has homology with other patients' CRKP. After taking the interventions, all the hospital beds and dining cars were pulled out to rinse with water, and the environment was disinfected.

**Supplementary Tables 1-8 are in separate Excel files and the table legends are as following:**

**Supplementary Table 1. Information of strains and patient diagnosis.**

**Supplementary Table 2. Phenotypes of 1566D, a.k.a., medical records of 1566P.**

**Supplementary Table 3. Phenotypes of 1567D, a.k.a., medical records of 1567P.**

**Supplementary Table 4. Phenotypes of 2035D, a.k.a., medical records of 2035P.**

**Supplementary Table 5. Phenotypes of 2036D, a.k.a., medical records of 2036P.**

**Supplementary Table 6. Phenotypes of 2037D, a.k.a., medical records of 2037P.**

**Supplementary Table 7. Phenotypes of 2038D, a.k.a., medical records of 2038P.**

**Supplementary Table 8. Phenotypes of 2039D, a.k.a., medical records of 2039P.**

#### Supplementary Table 9-12 are contained in the following of this document.

**Supplementary Table 9. Illumina MiSeq sequencing yields.**

| Sample | Total Reads | Raw Bases | Q20 Bases | Percentage (%) |
|---|---|---|---|---|
| 1566D | 3,743,220 | 1,125,453,384 | 941,700,320 | 83.67 |
| 2035D | 4,782,134 | 1,320,197,359 | 1,163,174,482 | 88.11 |
| 2036D | 5,856,910 | 1,483,756,319 | 1,311,194,394 | 88.37 |
| 2037D | 6,624,804 | 1,657,988,991 | 1,460,537,145 | 88.09 |
| 2038D | 5,615,282 | 1,597,685,382 | 1,368,045,328 | 85.63 |
| 2039D | 5,641,658 | 1,439,029,823 | 1,263,496,926 | 87.80 |
| 2040D | 4,801,236 | 1,220,424,147 | 1,075,399,273 | 88.12 |

**Supplementary Table 10. Oxford Nanopore sequencing yields.**

| Sample | Total reads | Total bases | Average length | Longest reads length | Reads N50 length |
|---|---|---|---|---|---|
| 1567D | 414,491 | 7,483,109,230 | 18,054 | 167,316 | 25,890 |

**Supplementary Table 11. Detection and validation of SNPs in seven strains.**

| Isolates | SNP number | cSNP number | cSNP percent (%) | Validated SNPs | Validation ratio (%) |
|---|---|---|---|---|---|
| 1566D | 8,499 | 7,336 | 86.3 | 154 | 77.0 |
| 2035D | 7,506 | 6,424 | 85.6 | 147 | 73.5 |
| 2036D | 33,716 | 28,835 | 85.5 | 152 | 76.0 |
| 2037D | 7,488 | 6,399 | 85.5 | 154 | 77.0 |
| 2038D | 8,734 | 7,497 | 85.8 | 156 | 78.0 |
| 2039D | 8,624 | 7,424 | 86.1 | 155 | 77.5 |
| 2040D | 8,880 | 7,636 | 86.0 | 152 | 76.0 |

**Supplementary Table 12. A total of 92 all-variation SNPs in seven strains.** Red refers to 40 all-variation loci; Bold stands for 24 strain's unique SNP loci.

| Chromsome | Position | Ref | 1566D | 2035D | 2036D | 2037D | 2038D | 2039D | 2040D |
|---|---|---|---|---|---|---|---|---|---|
| NC_016845.1 | 1066275 | T | C/T | C/T | T | C/T | C/T | C/T | C/T |
| NC_016845.1 | 1066289 | T | G/T | T/G | G/T | G/T | G/T | G/T | G/T |
| NC_016845.1 | 1157834 | G | G/T | T/G | G | G | G | G | G |
| NC_016845.1 | 1309634 | T | G | G | G | G | G | G | G |
| NC_016845.1 | 1309841 | A | T | A | A/T | T | T | T | T |
| NC_016845.1 | 1309847 | G | A | A/G | A | A | A | A | A |
| NC_016845.1 | 1309865 | G | A | A | A | A | A | A | A |
| NC_016845.1 | 1309889 | C | T | T | T | T | T | T | T |
| NC_016845.1 | 1311041 | G | C | C | C | C | C | C | C |
| NC_016845.1 | 1311056 | C | A | A | **C** | A | A | A | A |
| NC_016845.1 | 1311080 | G | G | C/G | C | C | C | C | C |
| NC_016845.1 | 1311134 | G | A | A | A | A | A | A | A |
| NC_016845.1 | 1311558 | G | T | T | T | T | T | T | T |
| NC_016845.1 | 1322931 | G | T | T | **G** | T | T | T | T |
| NC_016845.1 | 1322970 | A | G | G | G | G | G | G | G |
| NC_016845.1 | 1323036 | C | T | T | **C** | T | T | T | T |
| NC_016845.1 | 1324201 | A | C | C | C | C | C | C | C |
| NC_016845.1 | 1324306 | C | T | T | T | T | T | T | T |
| NC_016845.1 | 1324519 | G | A | A | A | A | A | A | A |
| NC_016845.1 | 1325195 | A | G | G | G | G | G | G | G |
| NC_016845.1 | 1325197 | C | T | **C** | T | T | T | T | T |
| NC_016845.1 | 1325233 | A | G | **A** | G | G | G | G | G |
| NC_016845.1 | 1325260 | A | A | A | A | C | C | C/A | C |
| NC_016845.1 | 3049023 | T | C | C | C | C | C | C | C |
| NC_016845.1 | 3060510 | T | C | C | C | C | C | C | C |
| NC_016845.1 | 3060570 | C | T | T | T | T | T | T | T |
| NC_016845.1 | 3060768 | C | G | G | G | G | G | G | G |
| NC_016845.1 | 3061023 | C | T | T | T | T | T | T | T |
| NC_016845.1 | 3074462 | C | T | T | T | T | T | T | T |
| NC_016845.1 | 3090625 | T | C | C | C | C | C | C | C |
| NC_016845.1 | 3090874 | A | G | G | G | G | G | G | G |
| NC_016845.1 | 3091363 | A | G | G | G | G | G | G | G |
| NC_016845.1 | 3097079 | G | G/A | G/A | G/A | G/A | G/A | G/A | G/A |
| NC_016845.1 | 3103775 | A | A/G | A/G | G | A | A/G | A | A |
| NC_016845.1 | 3103779 | T | T | T | **G** | T | T | T | T |
| NC_016845.1 | 3121888 | T | T | T | **C** | T | T | T | T |
| NC_016845.1 | 3121980 | A | A | A | **G** | A | A | A | A |
| NC_016845.1 | 3122814 | C | C/T | T/C | C/T | C | C | C | C |
| NC_016845.1 | 3122948 | A | A | A | **G** | A | A | A | A |
| NC_016845.1 | 3122951 | T | T/C | T/C | **C** | C/T | C/T | T/C | T/C |
| NC_016845.1 | 3123167 | G | G | G | **A** | G | G | G | G |
| NC_016845.1 | 3123235 | G | G | G | **A** | G | G | G | G |
| NC_016845.1 | 3123236 | A | A/T | A/T | T | A/T | A | A | A |
| NC_016845.1 | 3123266 | G | G | G | **A** | G | G | G | G |
| NC_016840.1 | 2673 | A | G | G | G | G | G | G | G |
| NC_016840.1 | 2677 | C | T | T | **T/C** | T | T | T | T |
| NC_016840.1 | 2707 | C | T | **T/C** | T | T | T | T | T |
| NC_016840.1 | 2737 | T | G | G | **G/T** | G | G | G | G |
| NC_016840.1 | 2812 | T | A | A | A | A | A | A | A |
| NC_016840.1 | 2896 | C | T | T | T | T | T | T | T |
| NC_016840.1 | 2905 | C | T/C | T | T/C | C/T | C/T | T/C | T/C |
| NC_016840.1 | 2932 | C | G | G | G | G | G | G | G |

| | | | | | | | | | |
|---|---|---|---|---|---|---|---|---|---|
| NC_016840.1 | 3013 | C | A | A | A | A | A | A | A |
| NC_016840.1 | 3070 | C | A | A | A | A | A | A | A |
| NC_016840.1 | 3094 | A | T | T | T | T | T | T | T |
| NC_016840.1 | 3120 | A | C | C | C | C | C | C | C |
| NC_016840.1 | 3223 | G | A | A | A | A | A | A | A |
| NC_016840.1 | 3244 | G | C | C | C | C | C | C | C |
| NC_016840.1 | 3250 | C | T/C | T/C | C/T | C/T | T | T | T |
| NC_016840.1 | 3299 | G | C | C | C | C | C | C | C |
| NC_016845.1 | 3578729 | A | **A/G** | A | A | A | A | A | A |
| NC_016845.1 | 3578759 | T | **T/C** | T | T | T | T | T | T |
| NC_016845.1 | 3578876 | T | C/T | T/C | T/C | C/T | C/T | T/C | T/C |
| NC_016845.1 | 3578930 | T | C/T | T | T/C | C/T | C/T | T | T |
| NC_016845.1 | 3579029 | T | T | T/G | G/T | G/T | T | T | T |
| NC_016845.1 | 3579152 | G | A/G | A/G | **A** | G/A | G/A | G/A | G/A |
| NC_016845.1 | 3579305 | T | T | T | **C/T** | T | T | T | T |
| NC_016845.1 | 3579389 | T | T/G | T/G | G/T | G/T | G/T | G/T | G/T |
| NC_016845.1 | 3579394 | G | G | G | **A** | G | G | G | G |
| NC_016845.1 | 3579425 | G | G/A | G | G/A | G | G | G | G |
| NC_016845.1 | 3579517 | T | T/C | T/C | T/C | C/T | C/T | T/C | T/C |
| NC_016845.1 | 4873352 | G | G | G | **G/A** | G | G | G | G |
| NC_016845.1 | 5169646 | A | A/G | G | A/G | G | G | G/A | G |
| NC_016845.1 | 5169731 | G | T/G | G/T | G/T | G/T | G/T | G/T | G/T |
| NC_016845.1 | 5169737 | A | A/C | C | C | C | A/C | C/A | C/A |
| NC_016845.1 | 5169743 | A | T/A | A/T | **T** | A/T | A/T | T/A | T/A |
| NC_016845.1 | 5169844 | G | T/G | G | T | G | G | G/T | G/T |
| NC_016845.1 | 5169852 | T | C | C | C | C | C | C | C |
| NC_016845.1 | 5169857 | T | C/T | C/T | T/C | C/T | T | T | T |
| NC_016845.1 | 5169860 | G | G/C | G/C | C | C | C/G | G/C | G/C |
| NC_016845.1 | 5169865 | G | T/G | G | G | G/T | G | G/T | G/T |
| NC_016845.1 | 5169881 | G | T/G | T | T | T | G/T | G/T | G/T |
| NC_016845.1 | 5169890 | C | C/A | A | A/C | A | C/A | C/A | C/A |
| NC_016845.1 | 5169900 | C | T/C | C/T | C/T | C/T | C/T | C/T | C/T |
| NC_016845.1 | 5169938 | A | A/G | A/G | A/G | A/G | A/G | A/G | A/G |
| NC_016845.1 | 5169950 | A | C/A | C/A | A/C | A/C | A/C | C/A | C/A |
| NC_016845.1 | 5169980 | A | C | C | C | C | A/C | C | C/A |
| NC_016845.1 | 5169989 | A | A/G | G | A/G | G | A/G | A/G | A/G |
| NC_016845.1 | 5169992 | C | C/G | G | C/G | G | C/G | G/C | G/C |
| NC_016845.1 | 5170059 | C | C/T | T | T/C | T | C/T | T/C | T/C |
| NC_016845.1 | 5170096 | G | G | C | G | C | G | G | G |
| NC_016845.1 | 5170099 | A | G/A | G/A | A/G | **G** | A/G | G/A | G/A |
